# Anamorphic development and extended parental care in a 520 million-year-old stem-group euarthropod from China

**DOI:** 10.1101/266122

**Authors:** Dongjing Fu, Javier Ortega-Hernández, Allison C. Daley, Xingliang Zhang, Degan Shu

## Abstract

Extended parental care (XPC) is a complex reproductive strategy in which progenitors actively look after their offspring up to – or beyond – the first juvenile stage in order to maximize their fitness. Although the euarthropod fossil record has produced several examples of brood-care, the appearance of XPC within this phylum remains poorly constrained given the scarcity of developmental data for Palaeozoic stem-group representatives that would link juvenile and adult forms in an ontogenetic sequence. Here, we describe the post-embryonic growth of *Fuxianhuia protensa* from the early Cambrian Chengjiang Lagerstätte, and show parental care in this stem-group euarthropod. We recognize fifteen distinct ontogenetic stages based on the number and shape of the trunk tergites, and their allocation between the morphologically distinct thorax and abdomen. Our data demonstrate anamorphic post-embryonic development in *F. protensa*, in which tergites were sequentially added from a posterior growth zone. A life assemblage consisting of a sexually mature *F. protensa* adult alongside four ontogenetically coeval juveniles, constitutes the oldest occurrence of XPC in the panarthropod fossil record. These findings provide the most phylogenetically basal evidence of anamorphosis in the evolutionary history of total-group Euarthropoda, and reveal a complex post-embryonic reproductive ecology for its early representatives.

## 1. Introduction

Fuxianhuiids comprise a distinctive clade of upper stem-group euarthropods exclusively known from the early Cambrian of South China [1-4], and figure among the most thoroughly scrutinized Lower Palaeozoic taxa owing to several remarkable instances of soft-tissue preservation [5-9]. Despite their contribution towards understanding the origin of Euarthropoda [3-10], most aspects of the post-embryonic development of fuxianhuiids and most other stem lineage representatives remain largely uncharted. Although it has been recognized that fuxianhuiid populations include individuals of different sizes and variable number of exoskeletal trunk tergites suggesting the occurrence of different ontogenetic stages [1-3, 5], there is no formal description of this variability nor its implications for the palaeobiology of these animals. Some studies have produced remarkable insights on the post-embryonic development in other Lower Palaeozoic stem-group euarthropods, such as the recognition of limb rudiments in a juvenile of the megacheiran *Leanchoilia illecebrosa* [11], and the changes in bivalved carapace morphology and body segment count during growth in *Isoxys auritus* [12]. However, detailed information on the ontogeny and reproduction of Lower Palaeozoic euarthropods is for the most part only available from crown-group members [13-22], precluding the ancestral reconstruction of these traits during the early evolution of the phylum. Here we describe the post-embryonic development of *Fuxianhuia protensa* from the Cambrian (Stage 3) Chengjiang biota in South China [1, 5]. This represents the most comprehensive characterization of the ontogeny in a stem-group euarthropod to date, and leads to the recognition of parental care in fuxianhuiids, casting new light on the complex reproductive behaviour of early animals during the Cambrian Explosion.

## 2. Materials and methods

The studied fossils are the result of decades of collecting effort from various localities of the Cambrian (Stage 3; locally Qiongzhusian) Yu’anshan Member of the Chiungchussu Formation of the Kunming region in South China [23]. The material is deposited in the Early Life Institute (ELI), Northwest University, Xian (see Supplementary Table S1). Specimens were prepared with fine needles under high magnification using stereomicroscopes. Fossils were photographed with a Canon EOS 5D Mark II digital camera and were processed in Adobe Photoshop CS 5. Camera lucida drawings were made using a Leica M80 microscope and prepared with Corel Draw X5. Morphometric data were measured from specimen photographs using the software FIJI [24].

## 3. Results

Complete individuals of *F. protensa* vary in total length from 1 to 8 cm (Fig. 1). All ontogenetic stages share a fundamentally similar body construction. The head comprises a (protocerebral) anterior sclerite with paired stalked eyes [25], a pair of pre-oral (deutocerebral) antennae [8], and a pair of para-oral (tritocerebral) specialized post-antennal appendages [2]. The anterior sclerite articulates with a subtrapezoidal head shield, whose proportions range from a 1:1 length/width ratio in juveniles (Fig. 1*a, b*; Fig. s1) to a wider 1:4 length/width ratio in later stages (Fig. 1*c-h*; Figs S2-S7). The trunk comprises a variable number of overlapping tergites and a terminal tailspine with paired caudal flukes. Although the trunk expresses most of the ontogenetic changes, there are some invariable aspects of its organization. 1) The head shield covers - but is not fused to - three reduced anteriormost tergites, each of which bears a single limb pair [1, 5]. 2) Trunk tergites gradually change in size posteriorly. 3) The trunk is morphologically divided into an anterior limb-bearing thorax with expanded tergopleurae, and a posterior limb-less abdomen composed of narrow cylindrical tergites. 4) Thoracic tergites are associated with two or three pairs of biramous walking legs as a result of a derived pattern of ventral segmental mismatch [1, 5, 26]. 5) The trunk endopods are largest in size and possess more podomeres towards the anterior half of the body across all ontogenetic stages, whereas they consistently have a reduced size and number of podomeres towards the posterior, with the smallest limbs located underneath the last thoracic tergite [1, 5] (Fig. 2; Fig. S1*a-f*).

**Figure 1.**
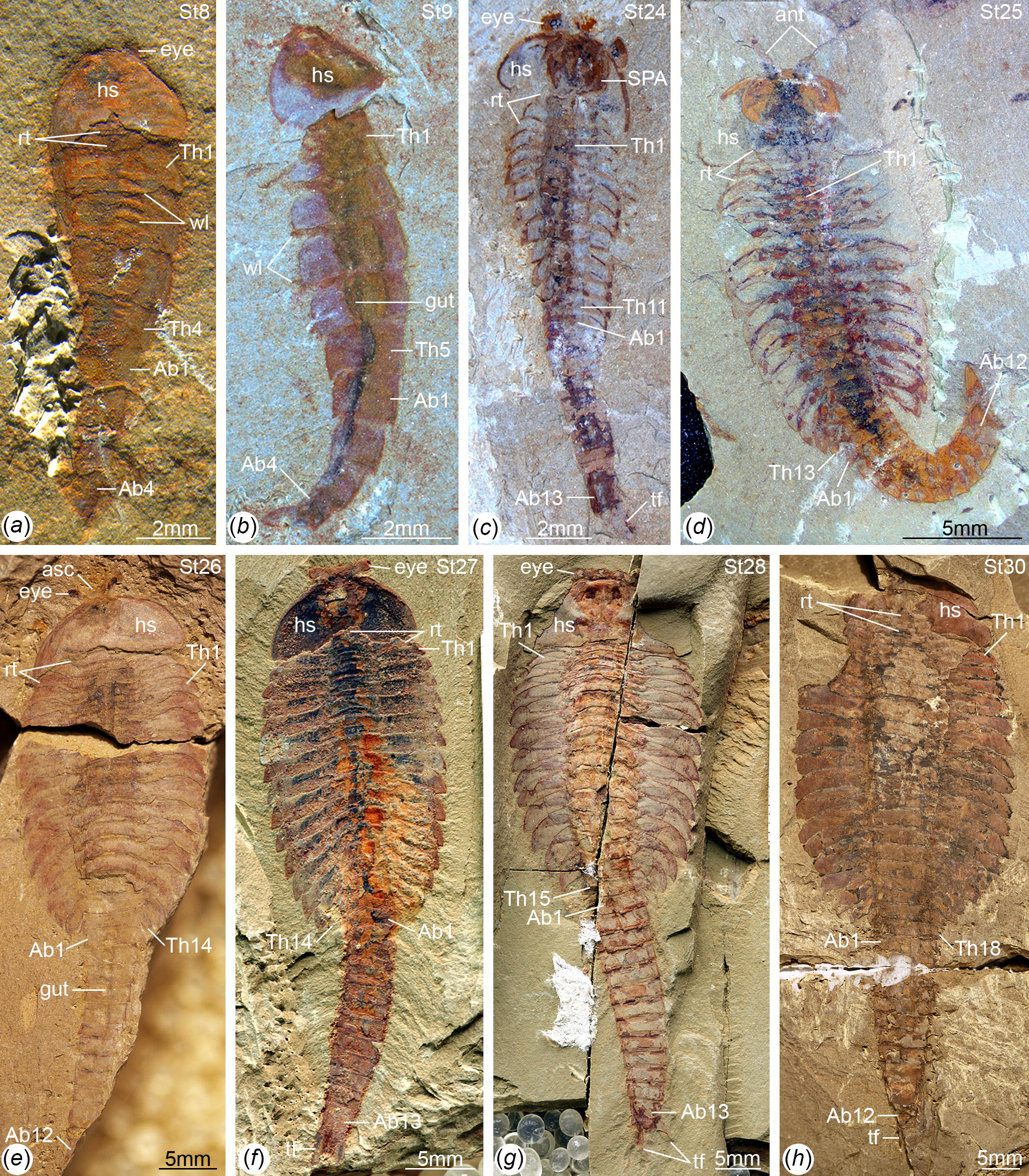
Ontogenetic stages of *Fuxianhuia protensa* from the early Cambrian Chengjiang Lagerstätte in South China. (*a*) ELI 0722A, stage 8; (*b*) ELI 0728, stage 9; (*c*) ELI 0034, stage 24a; (*d*) ELI 0050, stage 25b; (*e*) ELI 0001A, stage 26b; (*f*) ELI 36-77, stage 27a; (*g*) ELI 520-27A, stage 28a; (*h*) ELI 0011, stage 30b. Ab*n*, abdominal tergite; ant, antennae; asc, anterior sclerite; hs, head shield; rt, reduced anterior tergites; *tf*, tail flukes; Th*n,* thoracic tergite; *wl*, walking legs.

**Figure 2.**
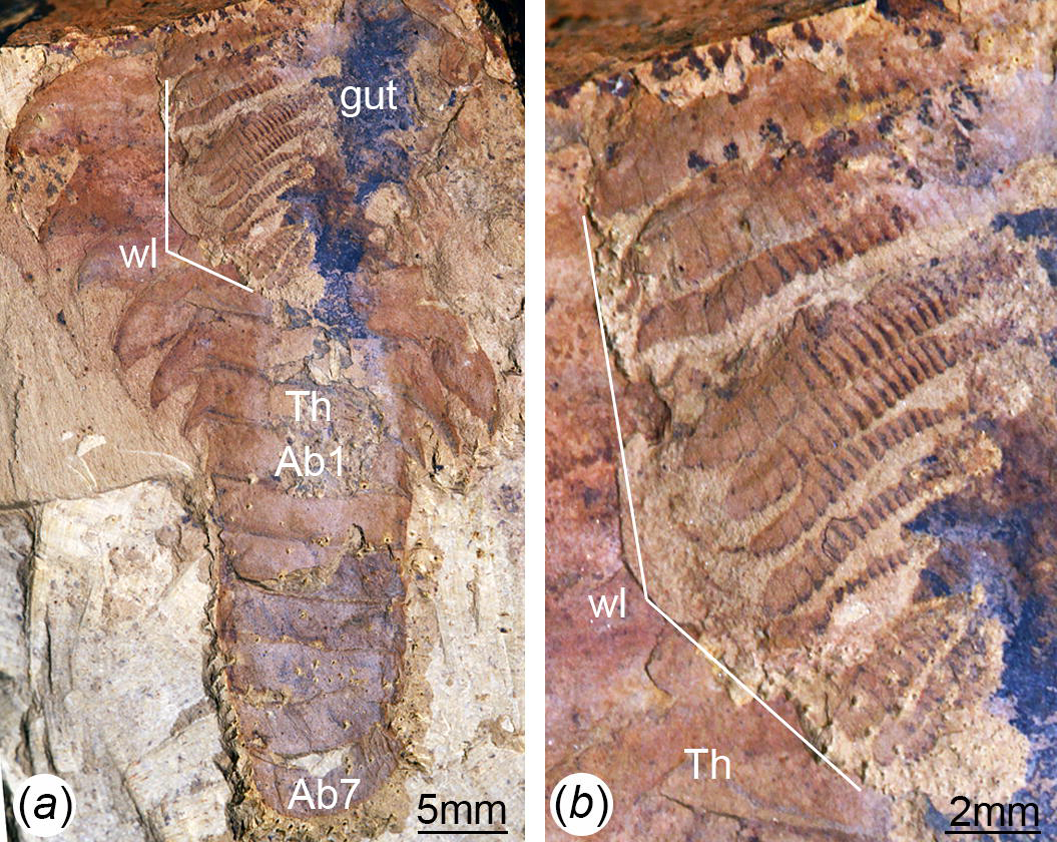
Trunk limb development in *Fuxianhuia protensa* from the early Cambrian Chengjiang Lagerstätte in South China. (*a*) CJ1069, articulated adult specimen of uncertain ontogenetic stage with dorsal exoskeleton prepared to illustrate the presence of multiple pairs of trunk appendages associated with each trunk tergite; note the morphological distinction between the limb-bearing thorax with expanded tergopleurae and the limb-less narrow abdomen. (*b*) Magnification of trunk appendages; note that endopods are less developed, smaller in size and have less podomeres towards the posterior end of the thorax. Ab*n*, abdominal tergite; Th*n,* thoracic tergite; *wl,* walking legs.

The available material allows us to directly recognize 15 distinct ontogenetic stages based on the number and shape of the trunk tergites, and their allocation between the thorax and abdomen (Fig. 1, 3; Figs s1-s8). In addition to the three reduced anteriormost tergites - which can be functionally considered as part of the head region despite the lack of cephalic fusion - complete individuals possess between 8 and 30 trunk tergites according to their degree of ontogenetic development (Fig. s9*a*). In stage 8 – the youngest juvenile available – the trunk consists of four limb-bearing thoracic tergites with short pleural spines (1:2 length/width ratio), and four limb-less abdominal tergites with a cylindrical outline (1:1 length/width ratio) (Fig. 1*a*, 3*a*,*b*; Fig. s1*a-j*). Thoracic tergites are approximately 1.5 times wider than those in the abdomen. Stage 9 is nearly identical to stage 8, differing only in the presence of five abdominal tergites (Fig. 1*b*; Fig. s1*k, i*). Individuals corresponding to stages 10 to 23 have not been recovered, but later ontogenetic phases demonstrate an increasing tergite count, and more substantial morphological differentiation. In stage 24, trunk tergites become broader and shorter, with the proportions being more pronounced in the tergites of the thorax (up to 1:6 length/width ratio) compared to those in the abdomen (1:2 length/width ratio) (Figs 1*c*, 3*a*; Fig. s2), and thoracic tergites being twice as wide as those in the abdomen. This phase provides insight into the transition from abdominal into thoracic tergites during ontogeny. Stage 24 individuals display either 11 thoracic and 13 abdominal tergites (see stage 24a in Fig. 3*a*; Fig. s1*a, b*), or 12 thoracic and 12 abdominal tergites (see stage 24b in Fig. 3*a*; Fig. s1*c, d*). The transformation of the oldest abdominal tergite into the youngest thoracic tergite is demarcated by the appearance of short tergopleural spines in the latter, which subsequently expand laterally throughout ontogeny (Figs 2 *a*, 3*a*), as well as the appearance of walking legs, which increase in size and number of podomeres towards the anterior end (Fig. 2*b*) [1, 5]. Likewise, the trunk tergites in stage 25 are allocated as either 12 thoracic and 13 abdominal (see stage 25a; Fig. s3*a-d*), or 13 thoracic and 12 abdominal (see stage 25b; Fig. s3*e-h*). These observations reveal a biphasic developmental pattern in which the trunk alternates between events of *accumulation* where a new tergite is added from the posterior growth zone resulting in an abdomen with 13 tergites, followed by *depletion* where the most anterior abdominal tergite becomes morphologically differentiated and is incorporated into the thorax, leaving the abdomen with 12 tergites (Fig. 3*a*). Critically, the thorax keeps incorporating new tergites with expanded tergopleurae one by one throughout ontogeny, increasing up to an observed maximum of 18 tergites (Fig. 1*h*, 3*b*; Figs s7-s9). This dynamic is maintained throughout later ontogeny (Fig. 1*d-h*; Figs s3-s7), with only rare individuals demonstrating a slight deviation of the pattern, such as the early integration of a thoracic tergite that results in a shortened abdomen with only 11 tergites (Fig. s3*i, j*). Stage 30 is the most mature developmental phase observed in our material, and is characterized by 18 thoracic and 12 abdominal tergites (Figs 1*h*, 3*a*; Figs s7-s9). Here, tergite proportions are even more pronounced (thoracic 1:13 length/width ratio; abdominal, 1:3 length/width ratio), and the thorax is up to four times wider than the abdomen. Taken together, the biphasic development and variability in the number of trunk tergites suggests that the complete ontogeny of *F. protensa* may have included up to 60 stages. This estimate carries the implication that the available material reflects approximately 25% of the post-embryonic development of this stem-group euarthropod, 20% of which is represented by advanced ontogenetic stages (Fig. s8).

**Figure 3.**
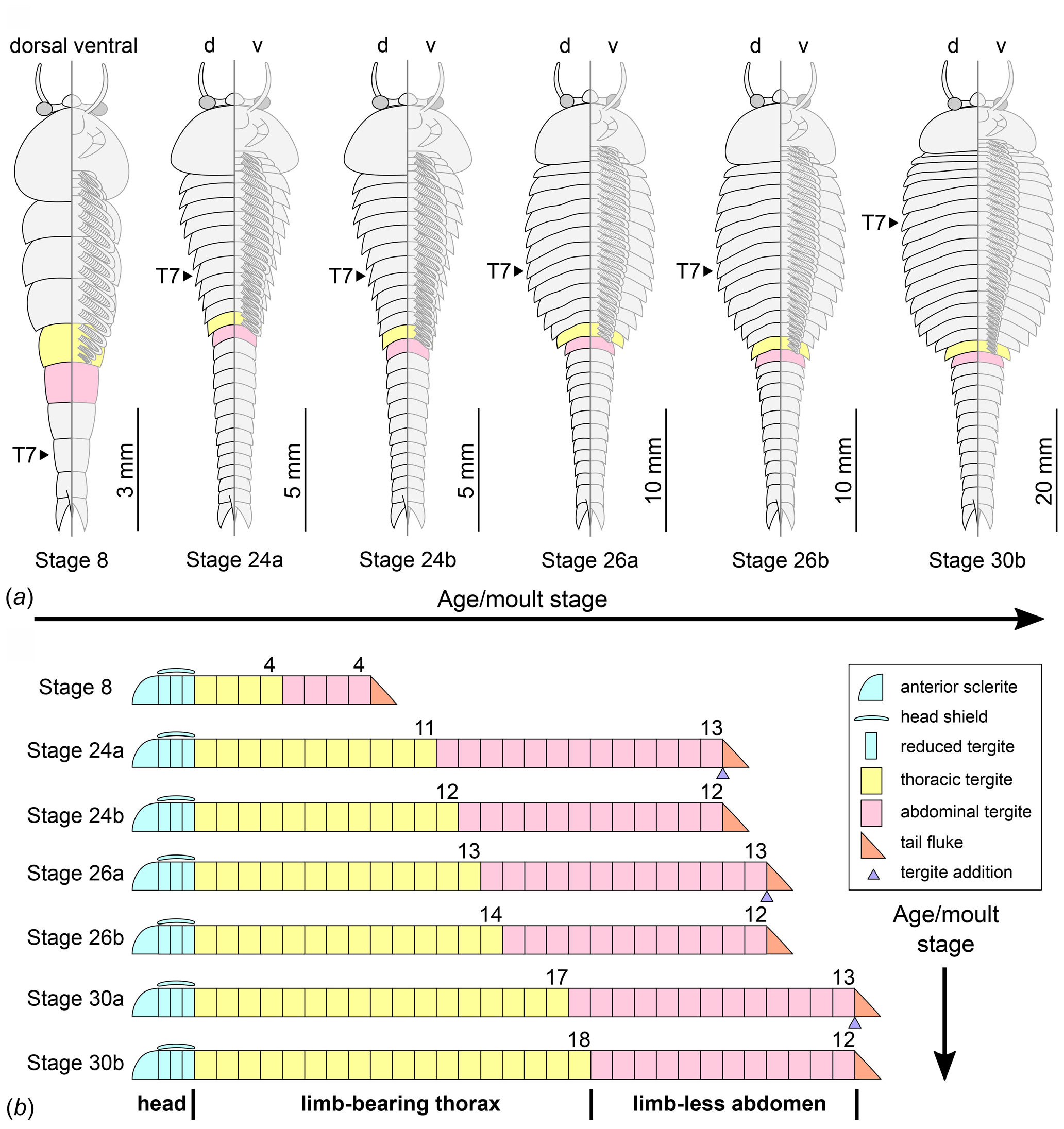
Ontogenetic changes in trunk region of *Fuxianhuia protensa* during anamorphic post-embryonic development. (*a*) Stage 8 is the earliest juvenile available, and possesses 4 thoracic limb-bearing tergites with short tergopleurae, followed by 4 limb-less cylindrical tergites. The three reduced anteriormost tergites under the head shield remain invariant. Throughout ontogenetic development, the most anterior abdominal tergite (pink) develops expanded pleurae and walking legs, transforming into the most posterior thoracic tergite (yellow). Tergite 7 arrowed in all stages for comparison. Stage 30b is the oldest known phase. (*b*) Partial trunk segmentation schedule for *F. protensa*; complete ontogenetic reconstruction provided in Fig. S8.

Morphometric data obtained from previously published and our newly documented specimens of *F. protensa* indicate that overall body size (estimated from thoracic length) is positively correlated with the number of trunk tergites, and that there is a substantial degree of body size variation among later ontogenetic stages (Fig. s9*b*). Since measurements were obtained from specimens collected from a number of different localities within the Yu’anshan Member of the Chiungchussu Formation (Supplementary Table S1), this pattern most likely reflects the natural size variation within these populations. Changes in the proportions of the dorsal exoskeleton also complicate morphometric measurements of overall body size. For example, some of the stage 8 or 9 individuals have a similar thoracic length to that of stage 24 specimens (Fig. s9*b*). This can be explained by the difference in trunk tergite proportions between instars, as stage 8 and 9 individuals have proportionately elongate thoracic tergites, whereas the thoracic tergites of Stage 24 are relatively shorter and wider.

The new ontogenetic data on *F. protensa* allows the interpretation of a complex fossil specimen that consists of one stage 29 individual preserved in close association with four stage 8 juveniles (Fig. 4*a-d*; Fig. s6*e, f*). All five individuals are extremely well preserved, and display a substantial degree of integrity as observed from the completely articulated dorsal exoskeletons, as well as the presence of delicate structures such as the stalked eyes, articulated head shield, trunk appendages, and gut tract. These observations strongly suggest that the five individuals were preserved in situ, with only negligible transport or pre-burial disturbance, and thus in all likelihood reflect a life assemblage rather than a time-averaged aggregation of random carcasses or exuviae.

**Figure 4.**
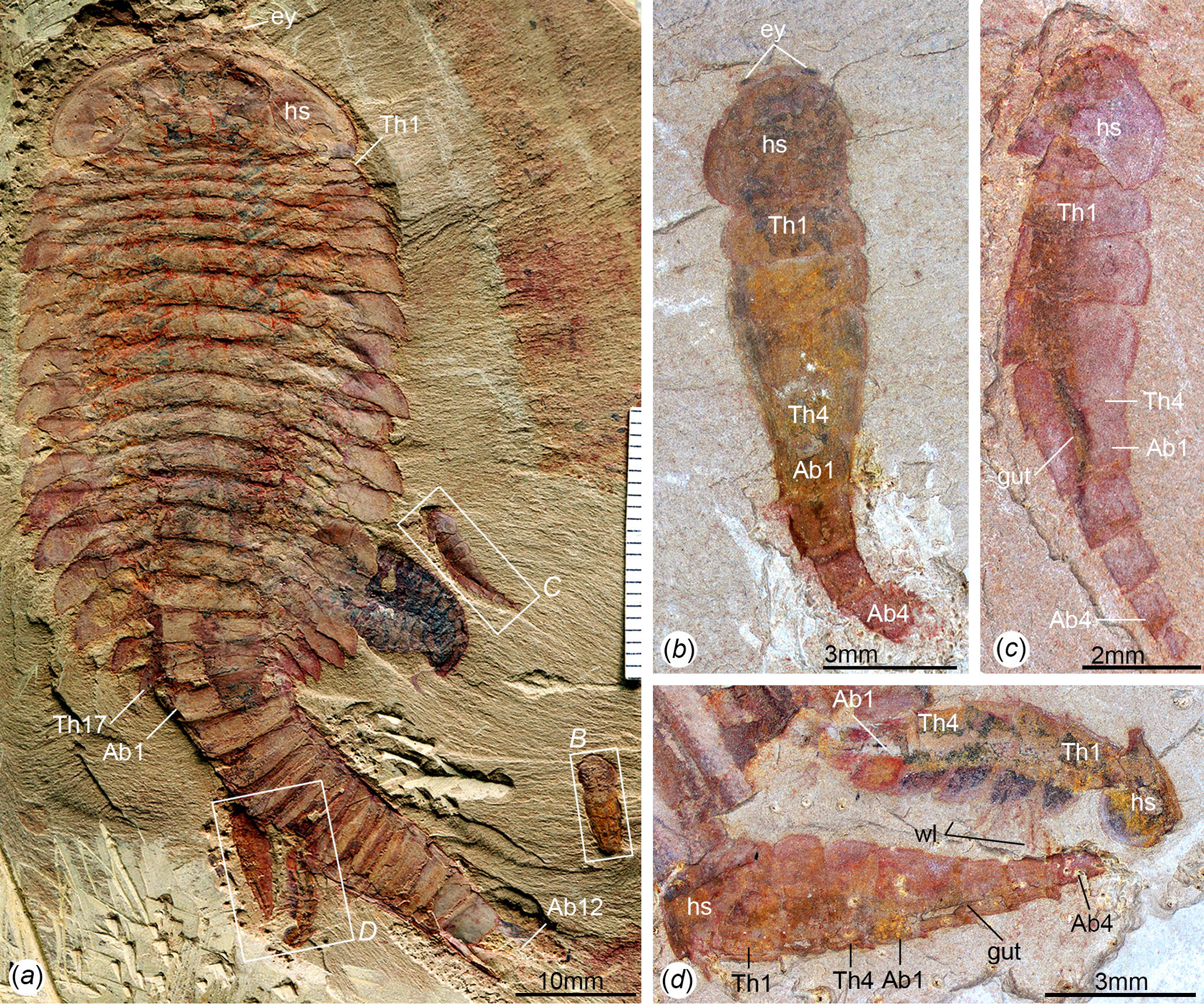
Fossil evidence for extended parental care in *Fuxianhuia protensa*. (*a*) ELI MU76A-a, life assemblage including a stage 29 – most likely sexually mature – adult individual and four stage 8 juveniles. (*b*) ELI MU76B-b, counterpart, articulated stage 8 juvenile with preserved eyes. (*c*) ELI MU76A-c, articulated stage 8 juvenile with preserved gut; note the presence of reduced tergites underneath head shield (see also Fig. S1). (*d*) ELI MU76A-d, two articulated stage 8 juveniles with preserved gut and walking legs. Abbreviations: Ab*n*, abdominal tergite; hs, head shield; tf, tail flukes; Th*n,* thoracic tergite; *wl*, walking legs.

## 4. Discussion

### (a) The ontogenetic development of *Fuxianhuia protensa*

Despite the gap in the ontogenetic series between stage 9 and stage 24 (Fig. s9), the smaller individuals can be reliably identified as representatives of *F. protensa* based on numerous shared morphological features. The presence of stalked eyes connected to an anterior sclerite, ventral antennae, a pair of specialized post-antennal appendages, limb polypody, and a loosely attached head shield that covers multiple reduced anterior tergites are collectively strong indicators of fuxianhuiid affinities (Fig. 1*a, b*; Fig. s1) [1-5]. The dorsal exoskeleton of stages 8 and 9 resembles *Chengjiangocaris* [1,2] and *Alacaris* [27] in the absence of a differentiated thoracic and abdominal regions; however, the possession of only three reduced anteriormost tergites underneath the head shield represents a key feature of *Fuxianhuia* [1, 2, 5, 27], whereas most other fuxianhuiid genera possess five [1, 2, 27] or six [3] reduced tergites. Although the presence of three reduced tergites is also known in *Guangweicaris spinatus* from the Guanshan biota (Cambrian Stage 4, Wulongqing Formation) [4], this taxon has never been reported from any of the stratigraphically older Chengjiang localities despite substantial collecting efforts on the Chiungchussu Formation [1, 5, 6, 8, 23, 26], and thus it is highly unlikely that the juvenile specimens are linked to it. In this context, the differences in the relative size and shape of the thoracic tergites between the *F. protensa* juveniles and adult specimens reflect ontogenetic change, as demonstrated by the gradual broadening of the thoracic tegites during stages 24 and 25 (Figs 1*a-d*; Figs s1-s3).

The ontogeny of *F. protensa* indicates that this taxon underwent anamorphic post-embryonic development (Fig. 3; Figs s8 and s9), in which new tergites were added sequentially from a posterior growth zone [17, 18, 22, 28, 29]. The biphasic developmental pattern of *F. protensa* is reminiscent of euarthropods that possess morphologically regionalized trunk regions, such as trilobites [22, 28 30] and some crustaceans [19, 31, 32]. Trilobite development is broadly characterized as hemianamorphic; it consists of an anamorphic phase in which new segments are added from a posterior growth zone after each moult (accumulation), followed by an epimorphic phase with an invariant number of segments that may involve the release of pygidial segments into the freely articulating thorax (depletion), or a body size increase without significant morphological changes [22, 28-30]. Although the mode of ontogenetic growth is somewhat variable between different trilobite taxa [22], the biphasic growth of *F. protensa* bears some broad similarities with that of *Shumardia* (*Conophyrs*) *salopiensis* [33]; both are typified by the alternation between posterior addition of segments, followed by the release of the anteriormost pygidial/abdominal segment into the thoracic region [22, 33]. However, the small sample size of the latest ontogenetic stages of *F. protensa* available (stage 30, *n* = 3; Supplementary Table S1) does not allow to unequivocally recognize an epimorphic phase (Figs s8, s9), nor to reliably distinguish the precise mode of anamorphic growth in this stem-group euarthropod (i.e. hemianamorphic, euanamorphic, teloanamorphic; see ref. 23].

The growth trajectory of *F. protensa* as recorded by the morphometric data (Fig. s9*b*) also evokes parallels with that of the Silurian proetid trilobite *Aulacopleura konincki*, in which body size is somewhat constrained among juveniles, but is more variable during late ontogeny resulting in adults with a polymorphic trunk segment count [34, 35]. The prevailing interpretation for the growth pattern of *A. konincki* favours a developmental model where the total number of thoracic segments was determined during early ontogeny, and once this maximum was reached, growth continued epimorphically [34, 35]. Unfortunately the small sample size of measurable juveniles (stage 8 and 9, *n* = 9; Supplementary Table S1) prevents us from drawing a comparable interpretation for the regulatory mechanisms responsible for *Fuxianhuia* development.

Among extant euarthropods, the biphasic development of the dorsal exoskeleton in *F. protensa* evokes comparison with that of Cephalocarida, a relatively poorly understood clade of meiofaunal crustaceans that possess a unique combination of ancestral and derived traits within the group [31, 32]. The post-cephalic exoskeleton of cephalocarids comprises a limb-bearing thorax with pivot-jointed tergites that possess laterally extended tergopleurae, followed by an elongate and flexible abdomen formed by ring-like tergites without appendages. During their early anamorphic development, the trunk of cephalocarids alternates between the production of new abdominal segments from a posterior growth-zone, and the transformation of the anteriormost abdominal tergite into a limb-bearing thoracic tergite with tergopleurae [31, 32]. Although there is some variation on the timing of these events between cephalocarid species [31], this overall pattern closely resembles the ontogenetic development of the trunk in *F. protensa* (Fig. 4; Figs s8, s9*a*). The development of Cephalocarida has been regarded as autapomorphic among crustaceans [31], which suggests that the growth mode of *F. protensa* most likely evolved independently given the phylogenetic position of Fuxianhuiida in the euarthropod stem lineage [27]. This interpretation is further supported by the lack of substantial dorsal exoskeletal tagmosis in closely related taxa such as *Chengjiangocaris* [1, 5], *Shankouia* [3] and *Alacaris* [27]. Ultimately, the ontogeny and phylogenetic position of *F. protensa* support anamorphosis as the ancestral mode of euarthropod post-embryonic development [11, 12, 17-20, 22], whereas the dorsal exoskeletal tagmosis of *Fuxianhuia* species [1, 2, 5] - and the closely related *Guangweicaris* [4] - are best regarded as a derived condition within upper stem-group Euarthropoda [10].

Our data also offer additional insights into the segmental organization of fuxianhuiids. It is well established that the thoracic tergites of *F. protensa* bear more than one set of biramous limbs, which raises the question of whether this type of segmental mismatch results from the derived organization of the dorsal or ventral side of the trunk [1, 2, 5, 9, 36]. The step-wise formation of the tergites from the posterior growth zone in the abdomen (Fig. 3; Figs s8, sS9), together with their release into the thoracic region and subsequent lateral expansion of the tergopleurae, suggest that the dorsal exoskeleton of fuxianhuiids follows the conventional pattern of segment production observed in most euarthropods [17-20, 28-31]. This condition suggests that the presence of multiple leg pairs associated with each thoracic tergite most likely represents a derived mode of ventral segmentation exclusive to this body region, which is further supported by the recent discovery of metamerically arranged midgut diverticulae in *F. protensa* that match the dorsal segmentation pattern [26]. Although rare, a similar type of dorsoventral segmental mismatch is also observed in the ventral trunk of the branchiopod *Triops cancriformis* [36, 37]. The anamorphic growth of *T. cancriformis* comprises the formation of limb-less abdominal segments from a posterior growth zone, which then develop between three and four pairs of limb buds that result in supernumerary thoracopods per segment in the adult [37]. This condition is closely reminiscent to that of *F. protensa*, including the fact that the newly formed posterior thoracic appendages are less developed compared to those on the anterior end of the body [37] (Fig. 2).

### (b) Extended parental care in *Fuxianhuia*

We interpret the exceptionally preserved in situ association of a stage 29 individual alongside four stage 8 juveniles as evidence of a parent and its offspring respectively (Fig. 4*a*, 5; Fig. s6*e, f*). Although it is not possible to pinpoint the exact growth stage at which fuxianhuiids became able to reproduce, the ontogeny of *F. protensa* identifies the stage 29 individual as developmentally advanced (Figs s8, s9), and thus in all likelihood a sexually mature adult. The fact that all the juveniles correspond to stage 8, and therefore have implicitly undergone a degree of post-embryonic development, identifies this association as a case of extended parental care (XPC), in which the parent (usually the female) nurtures the offspring during early ontogeny to maximize their survival and fitness [38-40]. This interpretation is supported by the fact that the four juveniles are ontogenetically contemporaneous, suggesting that they all belong to the same clutch. Unlike some cases of XPC in extant euarthropods or stratigraphically younger fossils, the adult *F. protensa* has no clear morphological adaptation for nursing the juveniles. For example, peracarid adult females have a specialized pouch for carrying the eggs and early juveniles [41, 42], and in some decapods the offspring may attach directly to the body of the parent [38, 39]. However, there are also recorded instances of XPC in which the parent cohabits with the juvenile for a prolonged period after hatching without the aid of specialized morphological adaptations [43]. In the absence of evidence for the presence of a brood pouch or other similar structure in *F. protensa*, we contend that this exceptional fossil association captures a case of prolonged cohabitation between parent and offspring [Fig. 4*e*, 5; Fig. s6*e, f*], not dissimilar from that observed in some extant marine crustaceans [38-40, 43]. Thus, *F. protensa* provides the phylogenetically and stratigraphically earliest evidence of XPC in the euarthropod fossil record [13-16, 41, 44], and reveals a greater diversity and complexity of reproductive strategies during the early Cambrian well beyond that of egg brood-care within a bivalved carapace [13-16].

**Figure 5.**
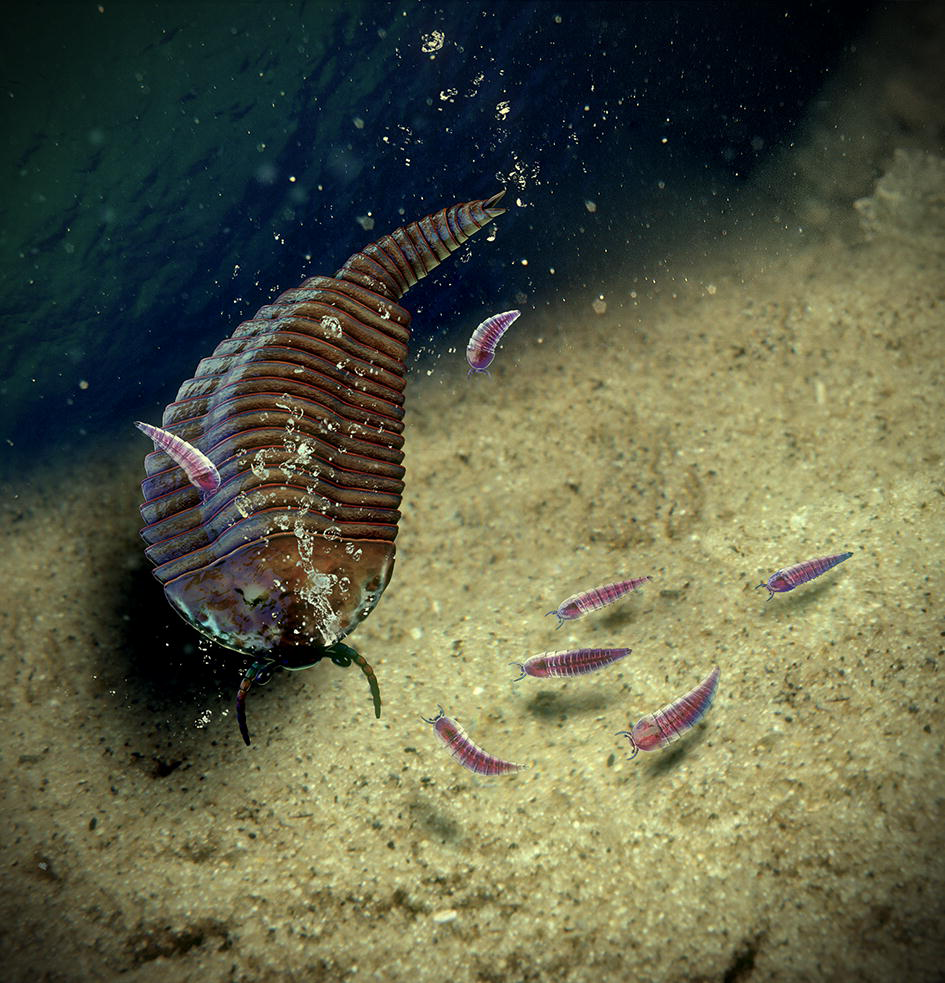
Reconstruction of extended parental care in *Fuxianhuia protensa*. Credit: Jingshan Fu.

## Data, Code, and Materials

The dataset and figures supporting this article have been uploaded as part of the supplementary material.

## Author contributions

D.-j.F., J.O.-H and A.C.D. performed research, wrote the paper and prepared the figures with input from the other authors. X.-l. Z., D.-g. S. and D.-j. F. designed the project with input of J.O.-H and A.C.D. All authors discussed and approved the manuscript.

## Acknowledgements

We thank Meirong Cheng, Juanping Zhai and Yu Wu at the Early Life Institute for help during fieldwork and technical assistance.

## References

1. Hou XG, Bergström J (1997) Arthropods of the Lower Cambrian Chengjiang fauna, southwest China. Fossils Strata 45: 1–116.

2. Yang J, Ortega-Hernández J, Butterfield NJ, Zhang XG (2013) Specialized appendages in fuxianhuiids and the head organization of early euarthropods. Nature 494: 468–471.

3. Waloszek D, Chen J, Maas A, Wang X (2005) Early Cambrian arthropods—new insights into arthropod head and structural evolution. Arthrop. Struc. Dev. 34: 189–205.

4. Yang J, Hou X, Dong W (2008) Restudy of *Guangweicaris* Luo, Fu et Hu, 2007 from the Lower Cambrian Canglangpu Formation in Kunming area. Acta Palaeontol. Sin. 47: 115–122.

5. Bergström J, Hou XG, Zhang XG, Clausen S (2008) A new view of the Cambrian arthropod *Fuxianhuia*. GFF 130: 189–201.

6. Ma X, Hou X, Edgecombe GD, Strausfeld NJ (2012) Complex brain and optic lobes in an early Cambrian arthropod. Nature 490: 258–261.

7. Ma X, Cong P, Hou X, Edgecombe GD, Strausfeld NJ (2014) An exceptionally preserved arthropod cardiovascular system from the early Cambrian. Nat. Comm. 5: 3560.

8. Ma X, Edgecombe GD, Hou X, Goral T, Strausfeld NJ (2015) Preservational pathways of corresponding brains of a Cambrian euarthropod. Curr. Biol. 25: 2969–297.

9. Yang J, Ortega-Hernández J, Butterfield NJ, Liu Y, Boyan GS, Hou JB, Lan T, Zhang XG (2016) Fuxianhuiid ventral nerve cord and early nervous system evolution in Panarthropoda. Proc. Nat. Ac. Sci. 113: 2988–2993.

10. Ortega-Hernández J (2016) Making sense of ‘lower’ and ‘upper’ stem group Euarthropoda, with comments on the strict use of the name Arthropoda von Siebold, 1848. Biol. Rev. 9: 255–273.

11. Liu Y, Melzer RR, Haug JT, Haug C, Briggs DEG, Hörnig MK, He YY, Hou XG (2016) Three-dimensionally preserved minute larva of a great-appendage arthropod from the early Cambrian Chengjiang biota. Proc. Nat. Ac. Sci. 113: 5542–5546.

12. Fu D, Zhang X, Budd GE, Liu W, Pan X (2014) Ontogeny and dimorphism of *Isoxysauritus* (Arthropoda) from the Early Cambrian Chengjiang biota, South China. Gond. Res. 25: 975–982.

13. Caron JB, Vannier J (2016) *Waptia* and the diversification of brood care in early arthropods. Curr. Biol. 26: 69–74.

14. Briggs DEG, Siveter DJ, Siveter DJ, Sutton MD, Legg D (2016) Tiny individuals attached to a new Silurian arthropod suggest a unique mode of brood care. Proc. Nat. Ac. Sci. 113: 4410–4415.

15. Siveter DJ, Tanaka G, Farrell ÚC, Martin MJ, Siveter DJ, Briggs, DEG (2014) Exceptionally preserved 450-million-year-old Ordovician ostracods with brood care. Curr. Biol. 24: 801–806.

16. Duan Y, Han J, Fu D, Zhang X, Yang X, Komiya T, Shu D (2014) Reproductive strategy of the bradoriid arthropod *Kunmingella douvillei* from the Lower Cambrian Chengjiang Lagerstätte, South China. Gond. Res. 25: 983–990.

17. Hughes NC, Haug JT, Waloszek D (2008) Basal euarthropod development: a fossil-based perspective. Evolving Pathways: Key Themes in Evolutionary Developmental Biology, 281–298.

18. Hughes NC (2007) The evolution of trilobite body patterning. Annu. Rev. Earth Planet. Sci. 35: 401–434.

19. Waloszek D, Maas A (2005) The evolutionary history of crustacean segmentation: a fossil-based perspective. Evol. Dev. 7: 515–527.

20. Walossek D (1993) The Upper Cambrian *Rehbachiella* and the phylogeny of Branchiopoda and Crustacea. Fossils Strata 32: 1–202.

21. Fortey RA, Hughes NC (1998) Brood pouches in trilobites. J. Paleontol. 72: 638–649.

22. Hughes NC, Minelli A, Fusco G (2006) The ontogeny of trilobite segmentation: a comparative approach. Palaeobiology 32: 602–627.

23. Hou XG, Siveter DJ, Siveter DJ, Aldridge RJ, Cong PY, Gabbott SE, Ma XY, Purnell MA, Williams M. (2017) The Cambrian Fossils of Chengjiang, China. The flowering of early animal life. Wiley Blackwell, 2nd Edition. 316 pp.

24. Schindelin, J. et al. (2012) Fiji: an open-source platform for biological-image analysis. Nat. Methods 9: 676–682.

25. Ortega-Hernández J (2015) Homology of head sclerites in Burgess Shale euarthropods. Curr. Biol. 25: 1625–1631.

26. Ortega-Hernández J, Fu DJ, Zhang XL, Shu DG (2018) Gut glands illuminate trunk segmentation in Cambrian fuxianhuiids. Curr. Biol. 28: R1–R2.

27. Yang J, Ortega-Hernández J, Legg DA, Lan T, Hou J, Zhang XG (2018) Early Cambrian fuxianhuiids from China reveal origin of the gnathobasic protopodite in euarthropods. Nat. Comms. 9: 470.

28. Hughes NC (2007) The evolution of trilobite body patterning. Annu. Rev. Earth Planet. Sci. 35: 401–434.

29. Minelli A, Fusco G (2013) Arthropod post-embryonic development. In Arthropod biology and evolution. Springer Berlin Heidelberg, p. 91–122.

30. Dai T, Zhang XL, Peng SC, Yao XY (2017). Intraspecific variation of trunk segmentation in the oryctocephalid trilobite *Duyunaspis duyunensis* from the Cambrian (Stage 4, Series 2) of South China. Lethaia 50: 527–539.

31. Olesen J, Haug JT, Maas A, Waloszek D (2011) External morphology of *Lightiella monniotae* (Crustacea, Cephalocarida) in the light of Cambrian ‘Orsten’ crustaceans. Arthrop. Struc. Dev. 40: 449–478.

32. Addis A, Biagi F, Floris A, Puddu E, Carcupino M (2007) Larval development of *Lightiella magdalenina* (Crustacea, Cephalocarida). Mar. Biol. 152: 733–744.

33. Stubblefield CJ (1926) Notes on the development of a trilobite, *Shumardia pusilla* (Sars). Zool. J. Linn. Soc. 36: 345–372.

34. Fusco G, Hughes NC, Webster M, Minelli A (2003) Exploring developmental modes in a fossil arthropod: growth and trunk segmentation of the trilobite *Aulacopleura konincki*. The American Naturalist 163: 167–183.

35. Hughes NC, Hong PS, Hou J, Fusco G (2017) The development of the Silurian trilobite *Aulacopleura koninckii* reconstructed by applying inferred growth and segmentation dynamics: a case study in paleo-evo-evo. Front. Ecol. Evol. 5: 37.

36. Ortega-Hernández J, Brena C (2012) Ancestral patterning of tergite formation in a centipede suggests derived mode of trunk segmentation in trilobites. Plos One 7: e52623.

37. Olesen J, Møller OS (2013) Notostraca. Atlas of Crustacean Larvae, eds Martin JW, Olesen J, Høeg T (John Hopkins Univ Press, Baltimore), pp. 40–46.

38. Thiel M (2000) Extended parental care behavior in crustaceans – a comparative overview. Crustacean Issues 12: 211–226.

39. Thiel M (2003) Extended parental care in crustaceans–an update. Revista Chilena de Historia Natural 76: 205–218.

40. Trumbo ST (2012) Patterns of parental care in invertebrates. The evolution of parental care. Oxford University Press, Oxford, 81–100.

41. Broly P, Serrano-Sánchez, MDL, Rodríguez-García S, Vega, FJ (2017) Fossil evidence of extended brood care in new Miocene Peracarida (Crustacea) from Mexico. J. Syst. Palaeontol. 15: 1037–1049.

42. Kobayashi T, Wada S, Mukai H (2002) Extended maternal care observed in *Parallorchestes ochotensis* (Amphipoda, Gammaridea, Talitroidea, Hyalidae). J. Crust. Biol. 22: 135–142.

43. Aoki M (1997) Comparative study of mother-young association in caprellid amphipods: is maternal care effective? J. Crust. Biol. 17: 447–458.

44. Wang B, Xia F, Wappler T, Simon E, Zhang H, Jarzembowski EA, Szwedo J (2015) Brood care in a 100-million-year-old scale insect. eLife 4: e05447.

